# Selective Peptidomimetic Inhibitors of NTMT1/2: Rational design, synthesis, characterization, and crystallographic studies

**DOI:** 10.1101/2020.04.13.040139

**Authors:** Brianna D. Mackie, Dongxing Chen, Guangping Dong, Cheng Dong, Haley Parker, Christine E. Schaner Tooley, Nicholas Noinaj, Jinrong Min, Rong Huang

## Abstract

Protein N-terminal methyltransferases (NTMTs) methylate the α-N-terminal amines of proteins starting with the canonical X-P-K/R motif. Genetic studies imply that NTMT1 regulates cell mitosis and DNA damage repair. Herein, we report the rational design and development of the first potent peptidomimetic inhibitors for NTMT1. Biochemical and co-crystallization studies manifest that **BM30** (IC_50_ of 0.89 ± 0.10 µM) is a competitive inhibitor to the peptide substrate and noncompetitive to the cofactor S-adenosylmethionine. **BM30** exhibits over 100-fold selectivity to NTMT1/2 among a panel of 41 methyltransferases, indicating the potential to achieve high selectivity when targeting the peptide substrate binding site of NTMT1/2. Its cell-permeable analog DC432 (IC_50_ of 54 ± 4 nM) decreases the N-terminal methylation level of SET protein in HCT116 cells. This proof-of principle study provides valuable probes for NTMT1/2 and highlights the opportunity to develop more cell-potent inhibitors to elucidate the function of NTMTs in future.

## INTRODUCTION

Protein methylation is an important epigenetic modification that regulates dynamic chromatin states and transcription^1,2^. Therefore, potent and specific inhibitors of protein methyltransferases can serve as valuable chemical probes to elucidate the function in physiological and pathological contexts. For example, recently approved Tazverik (tazemetostat) for epithelioid sarcoma treatment is an inhibitor for protein lysine methyltransferase (PKMT) EZH2. Another PKMT inhibitor, Pinometostat targeting DOT1L, is in a phase I trial for combination use for patients with acute myeloid leukemia ^1^. In addition, a few protein arginine methyltransferase 5 (PRMT5) inhibitors, including JNJ64619178, GSK3326595, PF06939999, and PRT811, are also in clinical trials for advanced solid tumors ^3^.

Protein α-N-terminal methylation has been known for nearly four decades since it was first uncovered on bacteria ribosomal proteins L33^4,5^. The identification of N-terminal methyltransferase 1 (NTMT1/NRMT1) unveiled the first methylation writer for human protein α-N-termini^6,7^. In addition to NTMT1, its close homolog NTMT2/NRMT2 that shares over 50% sequence similarity has been identified as another writer for human protein α-N-terminal methylation^8^. Both NTMT1 and 2 share the same X-P-K/R canonical recognition motif, where X can be any amino acid although D and E are not preferred^9–11^. Recently, METTL13 has been discovered to solely methylate the elongation factor 1A (EF1A) on both the α-N-terminus and lysine side chain^12,13^. However, the N-terminal sequence of EF1A is different from the X-P-K/R motif that NTMT1/2 prefer. Structural studies on NTMT1/2 in complex with their substrate peptides reveal that the peptide substrates bind at a defined binding pocket that juxtapose with the cofactor S-adenosyl methionine (SAM) binding site^9–11^. This is distinct from PKMTs and PRMTs, which normally have a narrow channel to accommodate only the side chain of the Lys or Arg. The structural information gained from the co-crystal structures substantiated the unique selectivity NTMT1/2^9–11^.

Protein α-N-terminal methylation was originally proposed to regulate protein-protein interactions since the methylated proteins initially identified were involved in large macromolecular complexes including histone proteins, cytochrome c-557, and myosin light chain proteins^4,14^. Recent discoveries have demonstrated its relevance in protein-DNA interactions, as shown in strengthening the interactions of chromatin with centromere protein A (CENPA) and regulator of chromosome condensation 1 (RCC1), as well as DNA damage-binding protein 2 (DDB2) to DNA damage foci^15–17^. Meanwhile, newly identified substrates of NTMT1 also include the tumor suppressor retinoblastoma 1 (RB1), oncoprotein SET, and poly(ADP-ribose) polymerase 3^6,18^. In addition, previous genetic studies have inferred the importance of NTMT1 in mitosis, DNA damage repair, aging, and a variety of cancers^6,16,17,19–21^. Despite the progress in recent years, our understanding of protein α-N-terminal methylation function is still in its infancy. Therefore, it is imperative to discover specific and potent inhibitors for NTMTs to probe their biological functions. Guided by the kinetic mechanism of NTMT1, bisubstrate analogs that covalently link both SAM and peptide substrate were designed to mimic the transition state and displayed high selectivity and potency for NTMT1^22–25^. Nevertheless, intrinsic properties of those bisubstrate analogs impose a challenge to penetrate cell membranes, which limit their applications in cell-based studies. Therefore, there is a need for a new strategy to discover inhibitors for NTMTs in order to probe the downstream process of protein α-N-terminal methylation. Given that the NTMT family has a unique peptide substrate binding site, we hypothesize that targeting the peptide binding site should achieve higher selectivity for NTMTs. Herein, we report the design and synthesis of the first potent and selective peptidomimetic inhibitors for the NTMT1/2.

## RESULTS AND DISCUSSION

### Design

According to the crystal structures of NTMT1 in complex with peptide substrates, the backbone carbonyl group of the first amino acid interacts with a conserved N168 residue NTMT1 through H-bonding^9,26^. Mutation of Asn168 to Lys resulted in a ∼36-fold loss in *K*_m_, indicating the importance of this interaction between NTMT1 and its peptide substrate^9^. However, NTMT1/2 was found to be able to methylate a variety of hexapeptides (XPKRIA) *in vitro*^11^. From the crystal structure, a conserved hydrogen bond is between the carbonyl oxygen of the first residue of the peptide substrate (X1) and the carboxamide side chain of Asn168^9^. The tolerance at the first position of the substrate suggests a possible spacious binding pocket surrounding the first position. The N-terminal amine of the peptide substrate points toward the methyl group of the SAM and serves as a nucleophile to receive the methyl group from the SAM. The second residue P2 forms a stacking interaction with the indole of Trp136. The third residue K3 forms two key hydrogen bonds with side chains of two aspartic acids^9–11^. Either a Lys or Arg residue has been discovered at the third position of known protein substrates. The fourth amino acid is adjacent to a negatively charged substrate binding channel, where the fifth and sixth residues are primarily exposed to the solvent^9–11^. Hence, we speculate that the α-N-terminal amine is essential for methyl transfer by acting as a nucleophile to attack SAM, but it is not essential for binding. Therefore, we hypothesized that the removal of the N-terminal amine of NTMT1/2 peptide substrates would convert the substrate into an inhibitor. Since the first four amino acids contribute significantly to the interaction with NTMT1/2, we initiated our efforts by incorporating different substitutions (**R**_**1**_) onto a tripeptide (PKR) through an amid bond (Figure 1) to retain as much interaction as the peptide substrates. PPKRIA and YPKRIA are two peptide substrates for NTMT1 with a *K*_d_ value at 0.7±0.2 nM and 0.1±0.08 µM, respectively^9^. Hence, compounds **1**-**5** were designed to mimic the Pro or Tyr moiety by introducing cyclopentane, cyclohexane, furan, phenyl, and a benzyl moiety as the **R**_**1**_ group.

**Figure 1.**
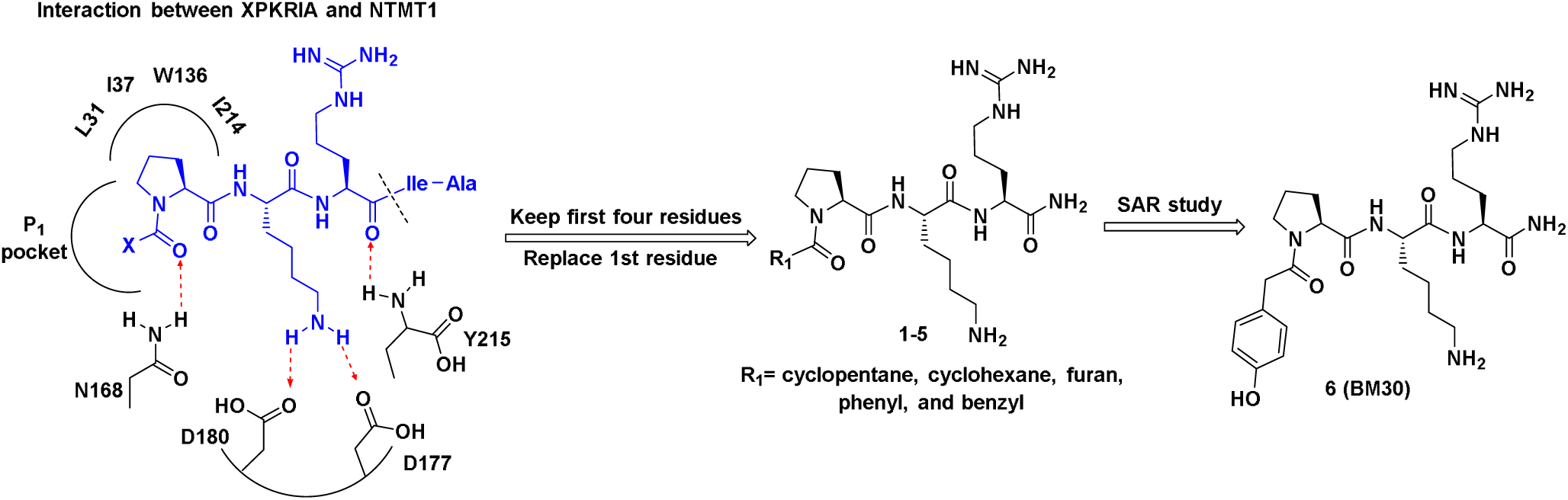
Design inhibitors based on the structure of NTMT1 peptide substrates. H-bond interactions are shown in red dotted lines.

### Synthesis

The peptides were synthesized following the standard Fmoc solid-phase synthesis on Rink amide resin using a CEM Liberty microwave peptide synthesizer. The peptidomimetics that have carboxylic acids at the first position were prepared through the standard amino acid coupling reaction (Scheme 1). All compounds were cleaved from the solid support in a cleavage cocktail consisting of trifluoroacetic acid/2,2’- (ethylenedioxy)diethanethiol/H_2_O/triisopropylsilane (94:2.5:2.5:1), and followed by purification through reverse-phase HPLC and characterized through mass spectrometry (Figure S1).

### Biochemical Characterization

All peptidomimetics were first subject to a methylation progression MALDI-MS study to examine if they were NTMT1 substrates^22,27^. None of the peptidomimetics displayed any methylation, indicating that they were not NTMT1 substrates. Next, the synthesized peptidomimetics were evaluated in a SAH hydrolase (SAHH)-coupled fluorescence assay at 100 µM of SAM and at the *K*_m_ value of the peptide substrate (SPKRIA or GPKRIA)^22^. Initial screening was carried out at the concentration of 5 µM, 30 µM, and 100 µM of each compound. Compound **5** exhibited more than 50% of inhibition at 30 µM, and was then subjected to subsequent IC_50_ studies by ranging the inhibitor concentration from 0.14 µM – 100 µM in a three-fold dilution. Compound **5** showed an IC_50_ of 6.85±2.82 µM (Figure S2A). A summary of the kinetic data for the peptidomimetics is shown in Table 1.

**Table 1.**
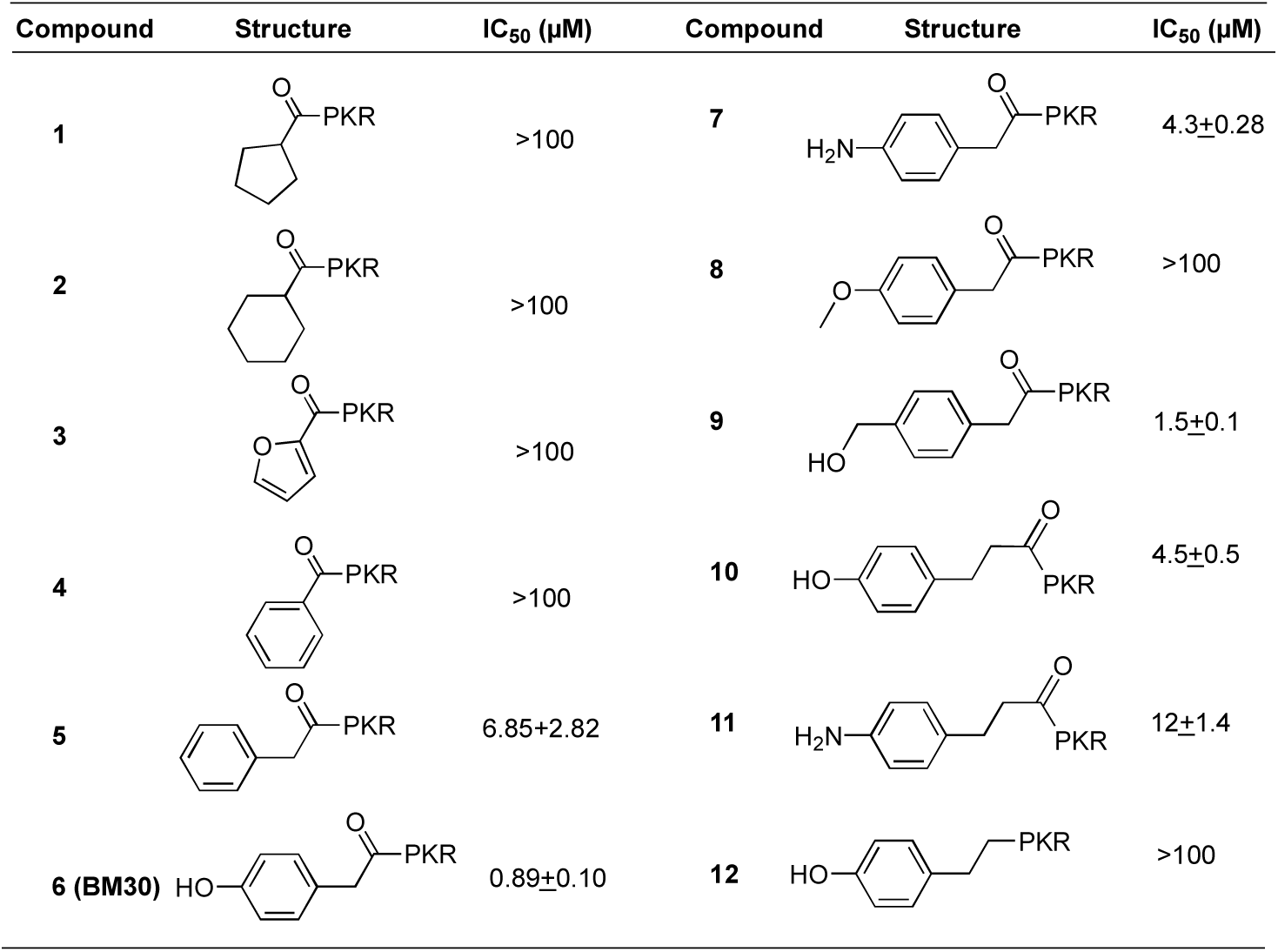
SAR of the first position of R_1_-PKR

| Compound | Structure | IC<sub>50</sub> (μM) | Compound | Structure | IC<sub>50</sub> (μM) |
| --- | --- | --- | --- | --- | --- |
| 1 |  | >100 | 7 |  | 4.3±0.28 |
| 2 |  | >100 | 8 |  | >100 |
| 3 |  | >100 | 9 |  | 1.5±0.1 |
| 4 |  | >100 | 10 |  | 4.5±0.5 |
| 5 |  | 6.85±2.82 | 11 |  | 12±1.4 |
| 6 (BM30) |  | 0.89±0.10 | 12 |  | >100 |

### Structure-Activity Relationship (SAR) Studies

In order to improve the inhibitory activity of the peptidomimetics, SAR studies were conducted by using **5** as the lead compound (Tables 1-2).

**Table 2.** SAR of the 2^nd^-4^th^ position of R_1_-PKR

| Compound | Structure | IC<sub>50</sub> (μM) | Compound | Structure | IC<sub>50</sub> (μM) |
| --- | --- | --- | --- | --- | --- |
| 13 |  | >100 | 19 |  | 4.8±0.71 |
| 14 |  | >100 | 20 |  | 13±1.6 |
| 15 |  | >100 | 21 |  | 16±4.7 |
| 16 |  | 21±3.3 | 22 |  | 4.2±1.2 |
| 17 |  | 1.0±0.22 | 23 |  | 6.4±0.5 |
| 18 |  | 4.8±0.31 | 24 |  | 32.9±8.3 |

**Table 3.** IC_50_ values and binding affinities of BM30 and its cell-permeable analogs

| Compound | Structure | IC<sub>50</sub> (µM) for NTMT1 | K<sub>d</sub> (µM) for NTMT1 | K<sub>d</sub> (µM) for NTMT2 |
| --- | --- | --- | --- | --- |
| BM30 |  | 0.89±0.10 | 3, 2.8 | 3.7 |
| DC431 |  | 0.23± 0.03 | 0.6, 0.8 | 3.4 |
| DC432 |  | 0.054± 0.004 | 0.3, 0.4 | 1 |

#### The first (R_1_) position

The benzoyl substitution of compound **5** resembled the side chain of Tyr; therefore, we focused our SAR investigation on introducing various para-substitutions on the phenyl ring, including a hydroxy, an amino, and a methoxy group at the para-position of the phenyl ring to yield compounds **6**-**8** (Table 2). Among all synthesized peptidomimetics, compound **6** (**BM30**) which contained a hydroxyl group at the para position showed the greatest inhibitory activity with an IC_50_ of 0.89±0.10 µM, approximately 7-fold increased activity compared to **5**. However, a methoxy substitution led to a loss of inhibition of 8 (IC_50_ > 100 µM), while its constitutional isomer **9** rescued the inhibitory activity (IC_50_ = 1.5±0.1 µM) with a 4-hydroxymethyl substitution. This suggests the preference of a H-bond donor at the first position. Given that introducing a methylene linker between the phenyl ring and the carbonyl group significantly increased the inhibition of **5** compared to **4**, an ethylene linker was introduced in **10** and **11** to probe the optimal linker length (Table 1). Compound **10** displayed a 5-fold loss in inhibition compared to **6**, while **11** exhibited a 3-fold loss in inhibition compared to **7**. Therefore, these results suggest that a methylene group is the optimal linker for the NTMT1 inhibitor. Lastly, compound **12** was prepared to explore the importance of the carbonyl group at the first position. Nevertheless, **12** did not show any inhibitory activity at 100 µM, implying the significance of the carbonyl group at the first position.

#### The second position

All available co-crystal structures of NTMT1/2 in complex with X-P-K/R substrates reveal that Pro2 only exhibits the trans conformation, suggesting that the trans conformation is favored to interact with NTMT1/2^9,11,26^. To verify the preference of this trans-conformation, an alpha-methyl-Pro was introduced at the 2nd position since the predominant conformation of alpha-methyl-Pro is cis.^28^ The resulting compound **13** completely lost the inhibitory activity for NTMT1 (Table 2). These results substantiate previous findings that the Pro2 prefers the trans conformation; however, the additional methyl group may also introduce certain steric hindrances. Next, ortho- and meta-amino benzoic acids were also introduced at Pro2 to generate **14** and **15**, respectively. Both compounds did not exhibit any inhibitory activity, denoting the importance of Pro2 in the canonical X-P-K/R motif.

#### The 3^rd^ and 4th position

Next, we explored the modifications at the third and fourth position in an attempt to increase inhibitory activity, as well as to enhance stability since peptide-based inhibitors are susceptible to degradation (Table 2). Arg has been detected at the 3rd position in the validated NTMT1 substrate CENP-A (GPRR)^10,15^, thus **16** was synthesized with Arg at the 3^rd^ position. However, **16** displayed over a 20-fold loss of inhibition compared to **6**. This result signified the preference for Lys over Arg at the third position, potentially due to its important role in interacting with two acidic residues (Asp177 and Asp180 in NTMT1; Asp232 and Asp 235 in NTMT2)^9–11^. Although Arg and Lys are two prevalent amino acids at the 4^th^ position of the NTMT1 substrate peptides, additional amino acids have been observed in reported NTMT1 substrates. Therefore, Lys, Thr, Ala, and Gly were introduced to replace the Arg at the 4^th^ position of **BM30** to generate **17**-**21**. Compound **17**, which included a Lys in the 4^th^ position, exhibited negligible difference in inhibition compared to **BM30** which contains an Arg. Introducing Thr or Ala at the 4^th^ position led to a 5-fold of loss as shown in **18**-**19**, while incorporating Gly produced almost a 14-fold loss in inhibition as shown in **20**. When the 4^th^ position was eliminated to yield **21**, inhibitory activity decreased 18-fold compared to **BM30**. Hence, the side chain of the 4^th^ amino acid does contribute to the NTMT1 interaction although it points towards to the solvent in the co-crystal structures. Additionally, other modifications were also made at the 4th position in attempts to increase the stability against proteases. Converting L-Arg to the non-natural amino acid D-Arg and beta-Arg in **22** and **23** resulted in approximately 4-fold and 6-fold loss of inhibition, respectively. Introducing a beta-Lys at the 3^rd^ position resulted in a 37-fold loss in activity, as shown in compound **24**.

### Selectivity Studies

Given that many methyltransferases exist, selectivity of the inhibitor is a critical requirement for its potential as a valuable probe. We first investigated the selectivity of **BM30, 7**, and **22** against one representative member from the protein lysine methyltransferase PKMT (G9a) and protein arginine methyltransferase PRMT (PRMT1) families. Both enzymes share a cofactor SAM binding site despite having different substrate peptide binding sites from NTMT1. As shown in Table S1, **BM30** and **7** demonstrated >100 µM IC_50_ values against G9a, while all three inhibitors displayed greater than >100 µM IC_50_’s against PRMT1. Thus, we proceeded with BM30 to evaluate its selectivity against a panel of 41 MTases (Reaction Biology) that include NTMT1/2, PKMTs, PRMTs, and DNMTs (Figure 2). The results indicated that BM30 displays comparable inhibition against both NTMT1 and NTMT2, which was not surprising since NTMT1/2 share the canonical X-P-K/R recognition motif. Strikingly, BM30 exhibited very marginal inhibition against the remaining 39 MTases even at 100 µM. The over 100-fold selectivity that BM30 displays for NTMT1/2 compared to PKMTs, PRMTs, and DNMTs evinces the unique substrate specificity of NTMT1/2.

**Figure 2.**
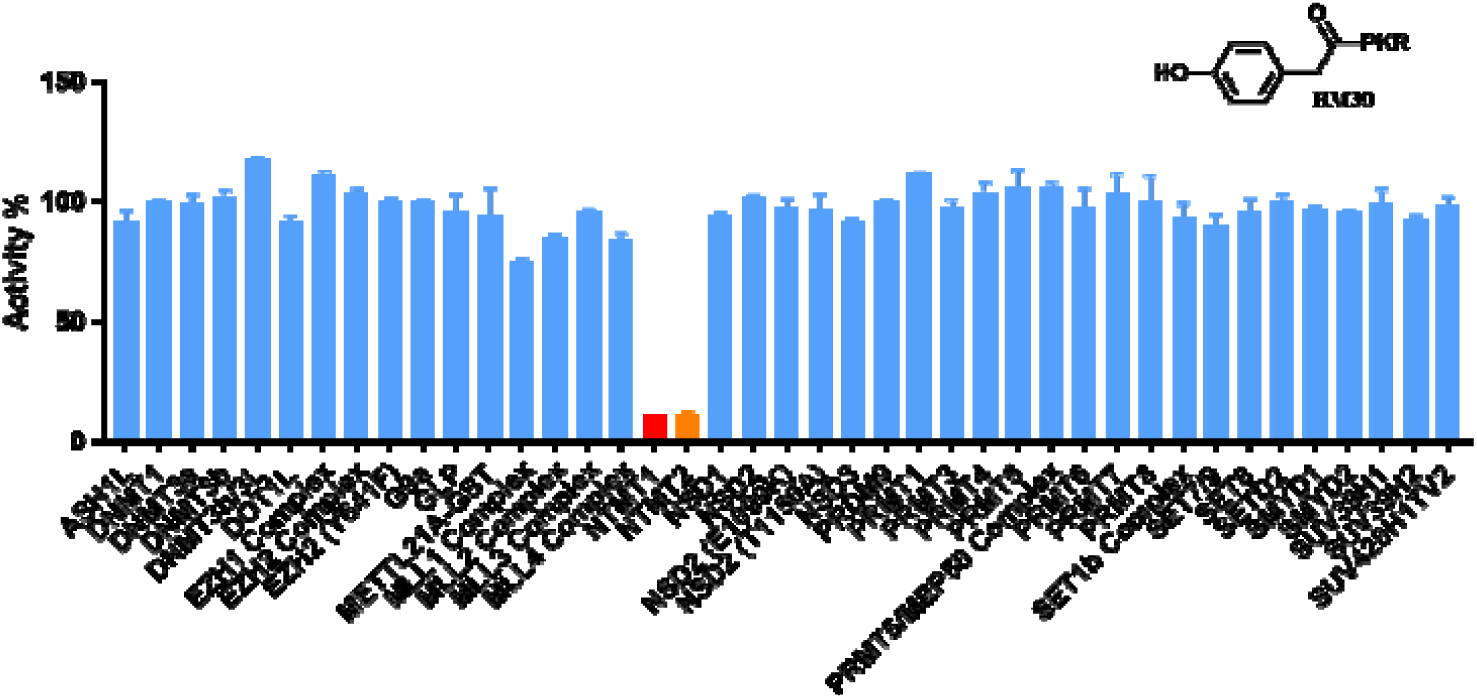
Selectivity of NTMT inhibitor **BM30** (100 µM for duplicate (n = 2)).

### Inhibition Mechanism

A kinetic analysis of **BM30** was performed to characterize its mechanism of actio using the SAHH-coupled fluorescence-based assay (Figure 3). As shown in Figure 3, **BM30** exhibited a competitive inhibition pattern with respect to the peptide substrate. This is demonstrated by an ascending and linear dependence of the IC_50_ values relative to the increasing peptide substrate concentration. On the other hand, **BM30** displayed a noncompetitive inhibition pattern with respect to the cofactor SAM, indicated by a line that is parallel to the x-axis. These results verify that **BM30** competitively binds to the peptide substrate binding site.

**Figure 3.**
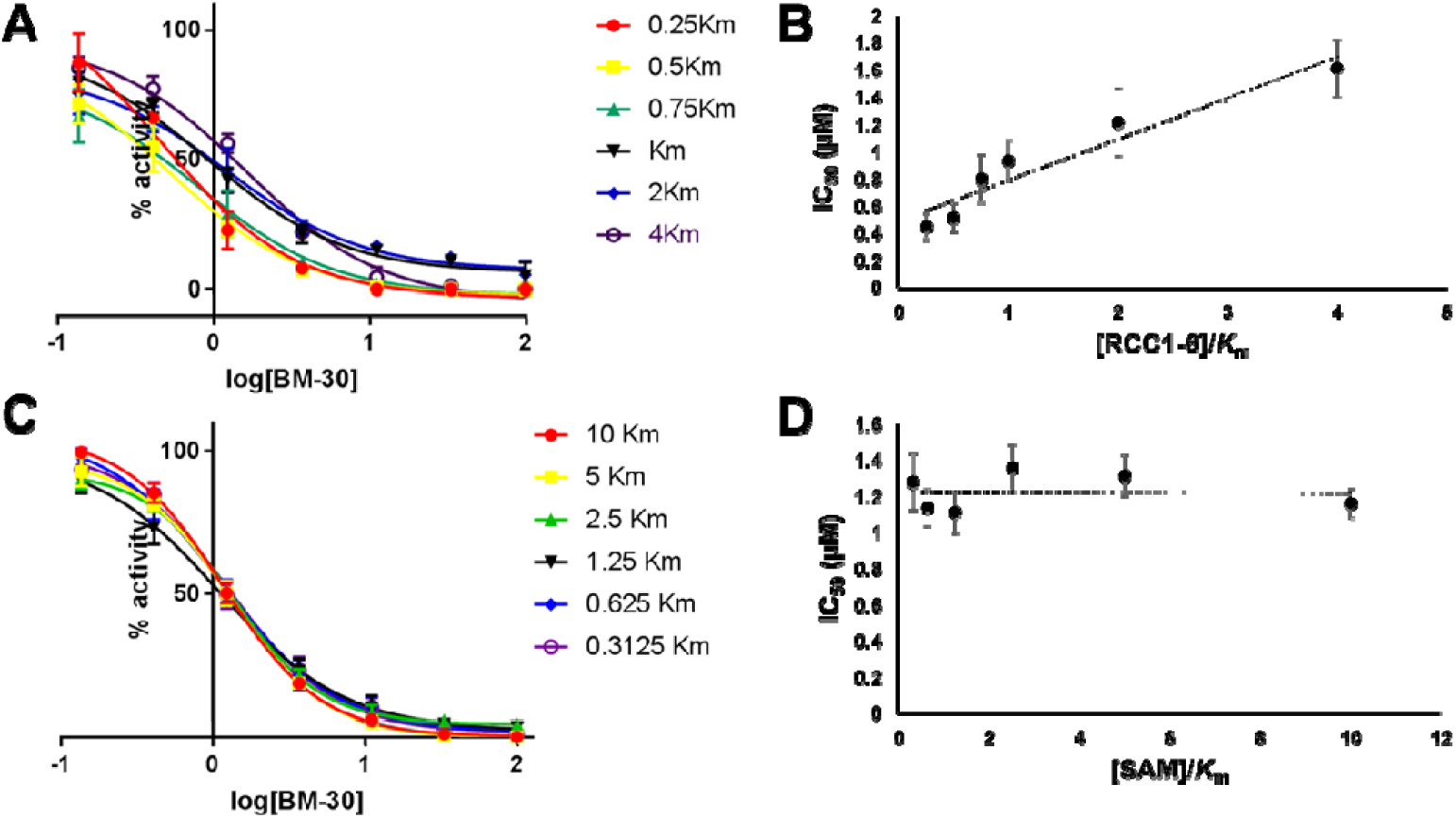
Inhibition mechanism of **BM-30**. (A) IC_50_ curves of **BM30** at varying concentrations of RCC1-6 with fixed concentration of SAM; (B) linear regression plot of IC_50_ values with corresponding concentrations of RCC1-6; (C) IC_50_ curves of **BM30** at varying concentrations of SAM with fixed concentration of RCC1-6; (D) Plot of IC_50_ values with corresponding concentrations of SAM.

### Crystal Structure of the NTMT1-BM30-SAH Complex

To further confirm that **BM30** binds to the peptide substrate binding site of NTMT1, we solved the X-ray co-crystal structure of NTMT1-**BM30**-SAH (PDB ID: 6WH8) (Figure 4A-C). **BM30** occupies the peptide substrate binding site of NTMT1. Superimposition of our NTMT1-**BM30**-SAH structure with the published NTMT1-YPKRIA-SAH ternary complex (PDB ID: 5E1D) gave an RMSD value of 0.12 Å (across all residues of chain A, Figure 4D-E). The **BM30**-NTMT1 complex retains the same interactions as previously observed with YPKRIA-NTMT1 in the ternary complex of substrate peptide/SAH^9^. For instance, the carbonyl oxygen of the 4-hydroxyl phenyl acetic acid interacts with the side chain of Asn168 through hydrogen bonding. The second Pro occupies a hydrophobic pocket that is formed by Leu31, Ile37, and Ile214. In addition, the ε-amine of the third Lys forms electronic interactions with carboxylate groups of Asp177 and Asp180. In addition, a direct H-bond exists between the carbonyl oxygen of the fourth Arg and Try215. The 4-hydroxyl group of the phenyl ring also forms a H-bond.

**Figure 4.**
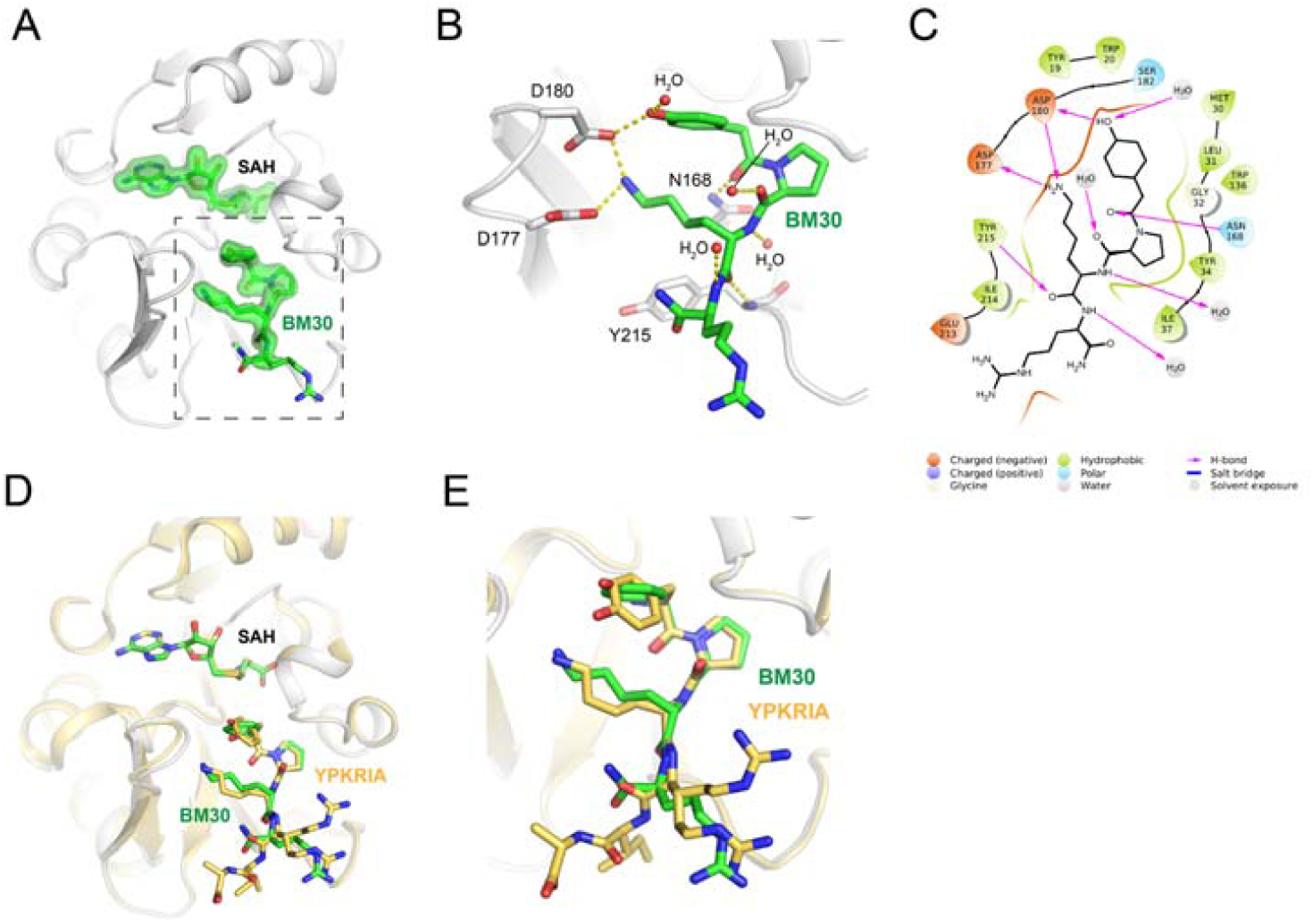
Co-crystal structure of NTMT1(gray cartoon)-**BM30** (green stick)-SAH (green stick) (PDB ID: 6WH8). (A) Fo-Fc omit density map of ligands. (B) Zoomed view of interactions of **BM30** with NTMT1. H-bond interactions are shown in yellow dotted lines. (C) **BM30** dinger Maestro) with NTMT1. (D) and (E) Structural alignment of NTMT1(gray)-**BM30** (green stick)-SAH (green stick) and NTMT1(yellow cartoon)-YPKRIA (yellow stick)-SAH (yellow stick) (PDB ID: 5E1D)

### Cell Permeable Analogs of BM30

We tested the cell permeability of **BM30** through MS detection of its cellular levels^29^. However, **BM30** was barely detected at 100 µM inside the cells (Figure S3). In order to increase the cell permeability, we designed and synthesized **DC431** and **DC432** by attaching cell-permeable peptides TAT and poly-Arg at the C-terminus of **BM30**, respectively. It is well-known that incorporating six Arg into compounds can increase the cell-permeability^30^. Hence, five additional Arg were added to **BM30** to generate **DC432** since **BM30** already had one Arg at its C-terminus. Given that the first four amino acids contribute significantly to the binding for NTMT1/2, we hypothesized that **DC431** and **DC432** would have slightly increased inhibitory activities than **BM30**, since they contained additional amino acids at the C-terminus of **BM30**. To confirm our hypothesis, IC_50_ values of **DC431** and **DC432** were determined to be 0.23±0.03 µM and 0.054±0.004 µM, respectively. **DC431** demonstrated a 4-fold increased inhibition for NTMT1 than **BM30**. Surprisingly, **DC432** displayed a 4-fold higher activity than **DC431** and 16-fold inhibitory activity compared to **BM30**. To further validate their interactions with NTMT1 and 2, we determined binding affinities of both cell-permeable analogs and **BM30** via isothermal titration calorimetry (Figure 5A-B). **BM30** exhibited a comparable *K*_d_ value of 2.9 µM and 3.7 µM to NTMT1 and 2, respectively. **DC431** showed a 4-fold increase in binding to NTMT1, although no improved binding to NTMT2 compared to BM30. On the other hand, **DC432** displayed an 8-fold tighter binding affinity to NTMT1, and a 4-fold tighter binding to NTMT2. Both **DC431** and **DC432** also exhibited enhanced cell permeability at 100 µM in comparison with the TAT peptide through a cellular MS assay (Figure S3).

**Figure 5.**
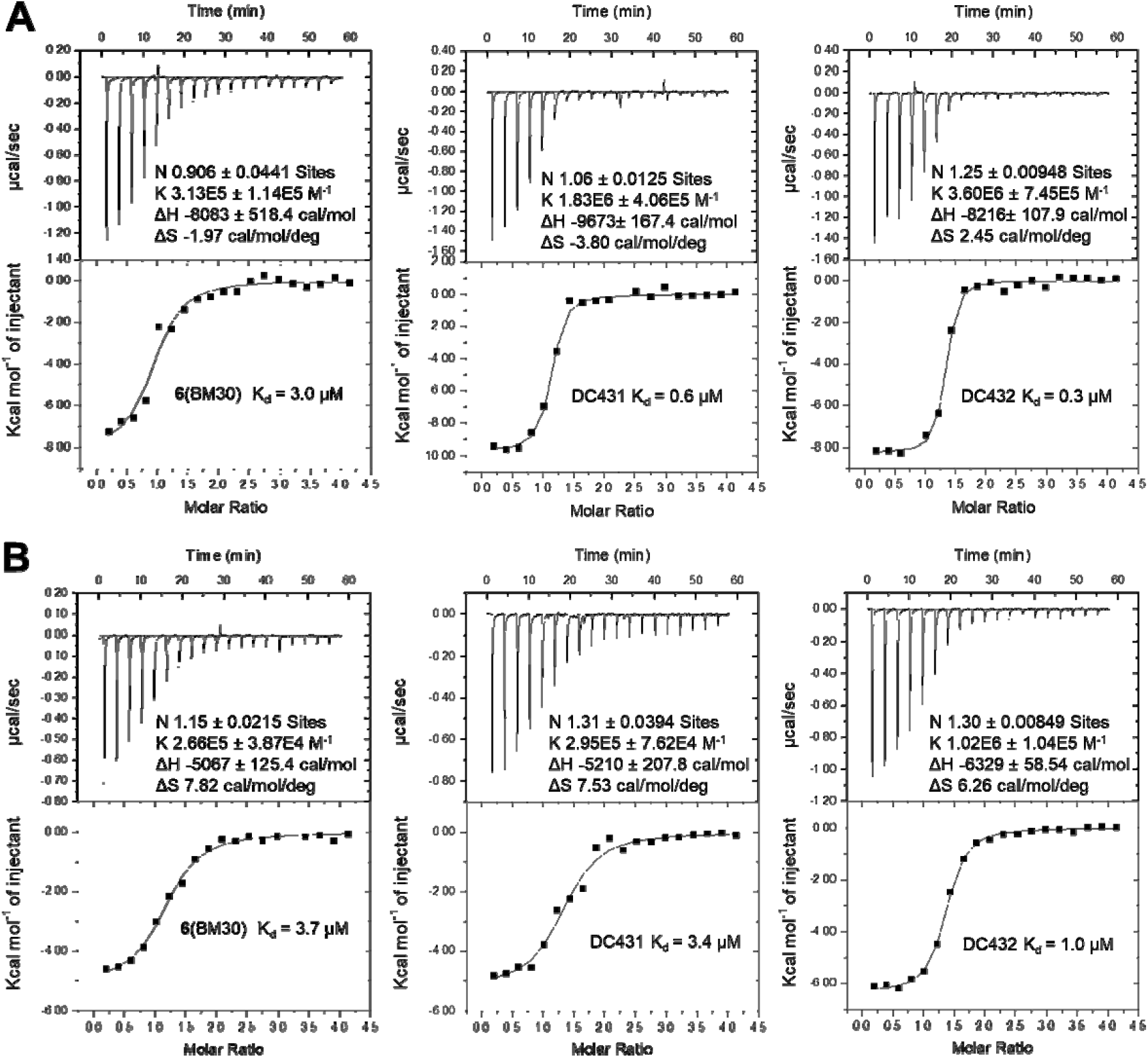
Binding affinity for NTMT1(A) and NTMT2 (B)

### N-terminal Methylation Level

NTMT1 can catalyze the trimethylation of its substrate starting with an SPK motif. Therefore, a cell permeable inhibitor of NTMT1 would be expected to decrease this trimethylation level. Then we proceeded to assess the inhibitory effects of **DC432** on the N-terminal trimethylation level via western blotting with a specific antibody to the N-terminal me3-SPK motif. As shown in Figure 6A, treatment with **DC432** for 72 h in HCT116 cells resulted in a decrease in the me3-SPK level (IC_50_ = 325 µM). This IC_50_ value was almost 1000-fold higher than the value obtained from the enzymatic assay. The discrepancy between these results may be due to the differences between peptide substrates and protein substrates or due to the limited cell permeability of **DC432**.

**Figure 6.**
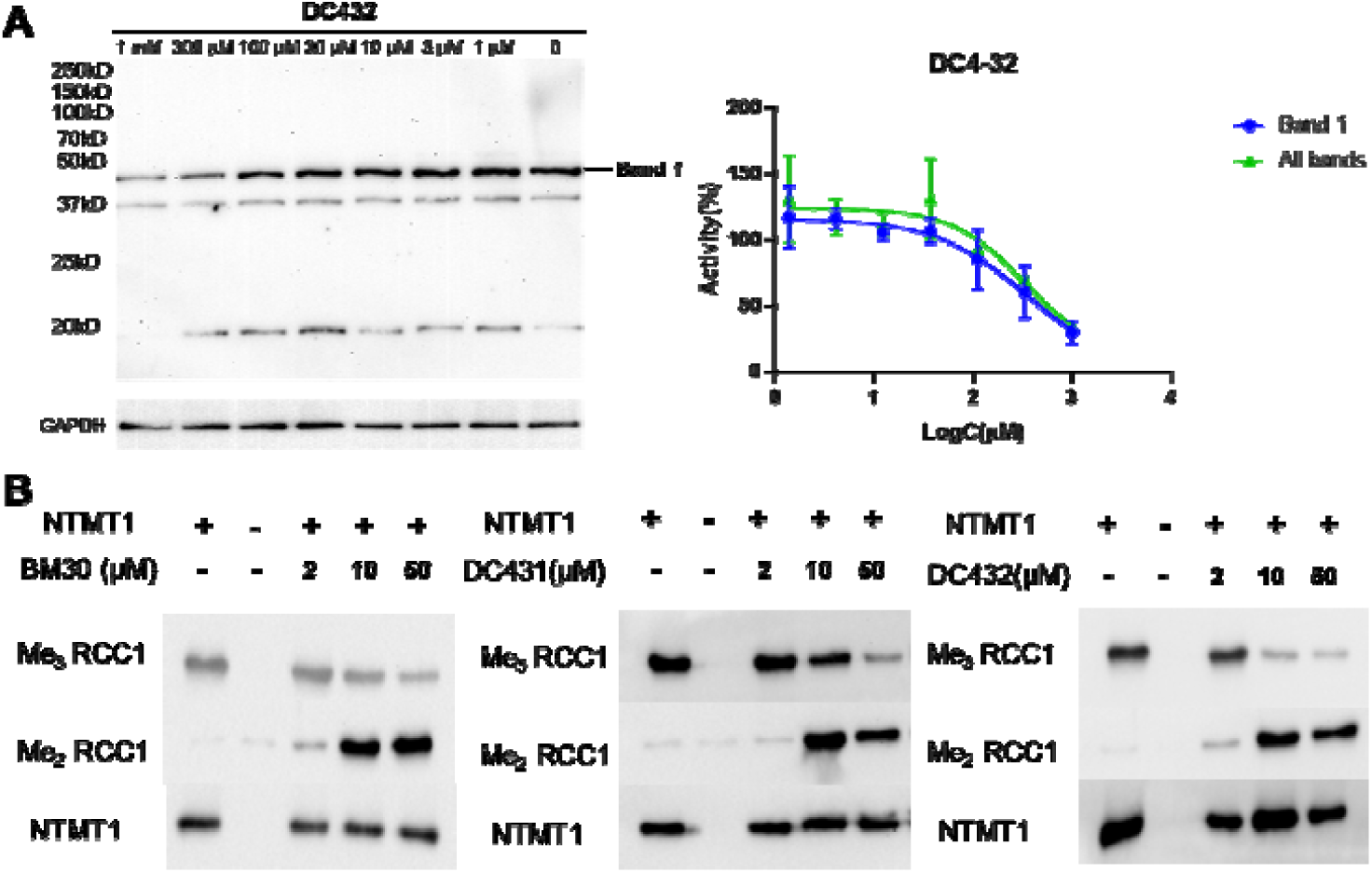
N-terminal methylation level assessment. (A) Cellular N-terminal methylation of compound DC432 in HCT116 cells. (B) N-terminal methylation of compounds BM30, DC431 and DC432 in the presence of NTMT1 in vitro

To further investigate these two possibilities, we tested the levels of N-terminal methylation by directly incubating the inhibitors with NTMT1, SAM and full-length RCC1 protein. At 2 µM of **BM30** or **DC432**, me2-RCC1 levels increased and me3-RCC1 levels decreased as compared to untreated samples (Figure 6B). The increased level of me2-RCC1 indicates inhibition of NTMT1 trimethylation, as NTMT1 is a distributive enzyme that normally reaches saturated trimethylation levels under these conditions^22,31^. The appearance and persistence of me2-RCC1 signals enzymatic inhibition. At 10 and 50 µM, all three inhibitors significantly decreased me3-RCC1 levels and increased me2-RCC1 levels (Figure 6B). These experiments with full-length RCC1 protein give an approximate IC_50_ value of 6 µM for **DC432**, which suggests the above discrepancy may be due to its limited cell permeability, as well as different inhibitory effects on peptide substrates from protein substrates.

## CONCLUSION

In summary, this study describes the discovery of the first potent and selective peptidomimetic inhibitor for NTMT1/2, **BM30**, reported to date. Furthermore, **BM30** selectively inhibits NTMT1/2 among a panel of 41 MTases. The cell-permeable derivative **DC432** represses N-terminal methylation levels of NTMT1/2 substrates in HCT116 cells, demonstrating that **DC432** would be a valuable probe for mechanistic studies of NTMT1/2 in cellular contexts. In addition, the crystal structure of the **BM30** in complex with NTMT1 revealed the structural basis for its potency and selectivity, which paves the way for the future design and development of additional cell-potent inhibitors.

## MATERIALS AND METHODS

### Chemistry General Information

All reagents and solvents were purchased from commercial sources (Fisher, Aldrich, and Chem-Impex) and used directly. Final compounds were purified on preparative high-pressure liquid chromatography (RP-HPLC) was performed on either Waters 1525 or Agilent 1260 Series system. Systems were run with 0-20% methanol/water gradient with 0.1% TFA. Matrix-assisted laser desorption ionization mass spectra (MALDI-MS) data were acquired in positive-ion mode using a Sciex 4800 MALDI TOF/TOF MS. The peptides (PKR, PKRIA and PRRRS) were synthesized on a CEM Liberty Blue Automated Microwave Peptide Synthesizer with the manufacturers standard coupling cycles at 0.1 mmol scale. All compounds were cleaved from the resin in a cocktail of trifluoroacetic acid/2,2’-(ethylenedioxy)diethanethiol/H_2_O/triisopropylsilane (94:2.5:2.5:1) and confirmed by mass spectrometry. The purity of final compounds was confirmed by Agilent 1260 Series system. Systems were run with 0-5% or 0-30% methanol/water gradient with 0.1% TFA. All the purity of target compounds showed >95%.

### NTMT1 Biochemical Assays

Kinetic characterization of the peptide inhibitors was determined in a fluorescence-based SAHH-coupled assay as described before. The inhibitors were added at concentrations ranging from 0 to 100 µM following a three-fold dilution. The methylation assay was performed under the following conditions: 25 mM Tris (pH = 7.5), 50 mM KCl, 0.01% Triton X-100, 5 µM SAHH, 0.2 µM NTMT1, 100 µM SAM, and 15 µM ThioGlo1. After 10 min of incubation at 37 °C, the reaction was initiated with 10 μL of 50 µM RCC1-6 for a total volume of 100 µL. Fluorescence intensity was monitored using a BMG ClarioStar microplate reader (Ex = 370 nm, Em=500 nm) at 37 °C for 15 min. The rates were fit to the log[inhibitor] vs response model using least squares nonlinear regression using GraphPad Prism 7 software. All experiments were performed in triplicate.

Inhibition Mechanism of BM-30 was performed using the fluorescent-based assay described above. Six independent IC_50_ studies of BM-30 were performed in triplicate with varying concentrations of substrate peptide, RCC1-6 and SAM at its *K*_m_ value. The inhibitors ranging in concentration (1 μL of 0.14 µM – 10 mM) and following a three-fold dilution were incubated in the well-solution that was added in the following order: ddH_2_O, buffer (10 µL of 250 mM Tris, pH 7.4 and 500 mM KCl), SAM (1 µL of 10 µM), SAH hydrolase (5 µL of 10 mg/mL), NTMT1 (0.5 µL of 40 µM) and ThioGlo1 (1 µL of 1.5 mM). After 10 min of incubation at 37 °C, the reaction was initiated with each concentration of RCC1-6 (10 μL of 5, 10, 15, 20, 40, 80 μM) for a total volume of 100 µL. The final concentrations are buffer (25 mM Tris, pH 7.4 and 50 mM KCl), SAM (1 µM), SAH hydrolase (10 µM), NTMT1 (0.2 µM), inhibitors (0 – 100 μM), RCC1-6 (0.25*K*m, 0.5*K*m, 0.75*K*m, *K*m, 2*K*m, and 4*K*m, 0.5, 1, 1.5, 2, 4, 8 μM, respectively) and Thioglo1 (15 µM). Fluorescence intensity was monitored using a CLARIOstar microplate reader (Ex = 370 nm, Em=500 nm) at 37 °C for 15 min. The rates were fit to the log[inhibitor] vs response model using least squares nonlinear regression through GraphPad Prism 7 software. The average IC_50_ value of each independent triplicate study was then plotted against the concentration of the [RCC1-6]/*K*m. Next, the same experiment was repeated with six IC_50_ studies of BM-30 in triplicate at varying concentrations of SAM and RCC1-6 at its *K*m value. The average IC_50_ value of each independent triplicate study were plotted against the concentration of the [SAM]/*K*m.

Selectivity Studies. To determine selectivity, kinetic analysis of the peptide inhibitors was initially carried out using the SAH hydrolase-coupled fluorescence assay against G9a and PRMT1, respectively. The inhibitors were added at concentrations ranging from 0 to 100 µM following a three-fold dilution. For G9a, the assay was performed in 1x PBS, pH7.4, 100 µM SAM, 10 µM SAH hydrolase, 25 nM G9a, inhibitors (0 – 100 μM), 20 μM H3-15 peptide, and 15 µM Thioglo1. For PRMT1, the assay was carried out in 2.5 mM HEPES (pH 7.0), 25 mM NaCl, 25 µM EDTA, 50 µM TCEP, 100 µM SAM, 10 µM SAH hydrolase, 200 nM PRMT1, 100 μM H4-12 peptide, and 15 µM Thioglo1. Fluorescence intensity was monitored using a CLARIOstar microplate reader (Ex = 370 nm, Em=500 nm) at 37 °C for 15 min. The rates were fit to the log[inhibitor] vs response model using least squares nonlinear regression through GraphPad Prism 7 software. All experiments were performed in triplicate.

### Isothermal Titration Calorimetry

Isothermal titration calorimetry (ITC) measurements were performed at 25 °C by VP-ITC MicroCal calorimeter. Purified NTMT1 and NTMT2 were diluted with the ITC buffer (20 mM Tris-HCl pH 7.5, 150 mM NaCl and 0.5 mM TECP) to the final concentration of 30-50 μM, and the compounds were dissolved to 0.5-1mM in the same buffer. The compound was titrated into the protein solution with 26 injections of 10 μl each. Injections were spaced 180 sec with a reference power of 15 μcal/sec. The ITC data were processed with Origin 7.0 software (Microcal).

### Co-crystallization and Structure Determination

Purified NTMT1 at 30mg/ml was mixed with BM-30 at a molar ratio of 1:2, incubated for 1 hr at 4°C. Broad matrix crystallization screening was performed using a Mosquito-LCP high throughput crystallization robot (TTP LabTech) using hanging-drop vapor diffusion method at 20°C. Crystals containing BM-30 were grown in 0.1 M Na-HEPES, pH 7.5, 1.6 M ammonium sulfate, and 2% PEG 1000; all crystals were harvested directly from the drop and flash cooled into liquid nitrogen. Data were collected on single crystals at 12.0 keV at the GM/CA-CAT ID-D beamline at the Advanced Photon Source, Argonne National Laboratory. The data were processed using the on-site automated pipeline with Fast DP and molecular replacement performed with Phaser-MR (PHENIX).^31-32^ All model building was performed using COOT (79) and subsequent refinement done using phenix.refine (PHENIX).^32-33^ The structure has been solved, refined and deposited into the Protein Data Bank with ID: 6WH8. Structure-related figures were made with PyMOL (Schrödinger) and annotated and finalized with Adobe Photoshop and Illustrator.

### Cell Permeability Evaluation by MALDI

The colon cancer cell line HCT116 was cultured in Mccoy’s media supplemented with 10% fetal bovine serum and 1% Antibiotic-Antimycotic (Gibco). The cells were cultured in tissue culture dish (Falcon 353003). Cells were maintained in cell culture flasks until seeding into a 12 well tissue culture plate (Falcon 353047). Media was removed and the cells were washed with DPBS (1 mL) twice followed by treating with TrypLE Express (1 mL) into a 100 × 20 mm culture flask. The reaction was quenched by addition of 4 mL of media and the cells were counted. Cells were seeded into a 12 well tissue culture plate (Falcon 353047) at a density of 0.1 × 10^6^ cells/mL and incubated overnight at 37 °C, 5% CO_2_, and 95% humidity, with the lid on. They were then treated with the inhibitors at different concentrations and incubation was continued for the specified time. TAT peptide (GRKKRRQRRR-NH_2_) was also incubated as a control experiment. The media was removed, and the cells were washed with 1x PBS three times to remove any residual compound or peptide attached to the cell surface. Then 100 µL of 1 x PBS was added and the cells were snap freeze in liquid nitrogen twice. The cell lysate was then analyzed with MALDI using DHB matrix to identify the presence of the compound inside the cell.

### Cellular N-methylation Level

HCT116 cells were seeded 20,000 cells/well on 24-well plates in the presence of 1x PBS or DC4-32 at different concentrations. The cells were harvested at 72 h in 1x RIPA cell lysis buffer (25mM Tris-HCl pH 7.6, 150 mM NaCl, 1% NP-40, 1% sodium deoxycholate, 0.1% SDS, protease inhibitors) and incubated for 30 min on ice. The cell lysates were centrifuged at 15000 rpm for 10 min and the precipitates were removed. The concentration of total protein was quantified by bicinchoninic acid (BCA) protein assay kit (ThermoFisher, #23228). Equal amounts of total protein were mixed with 4x loading dye and loaded onto a 12.5% SDS–PAGE gel and separated. The gel was transferred to a polyvinylidene difluoride (PVDF) membrane using Biorad Trans-Blot Turbo system. The membrane was then blocked for 1□h in 5% milk TBST solution and washed with 1x TBST solution three times. The membrane was incubated with anti-me3-SPK antibody at 4□°C for 12□h, washed with 1x TBST solution three times and then incubated with Rabbit IgG-HRP antibody (Cell Signaling, #7074S) for 1□h at room temperature. The membrane was again washed with 1x TBST solution three times and detected using a Protein Simple FluorChem imaging system. The data was analyzed in GraphPad prism software.

### *In Vitro* Methylation Assays with Full-length Recombinant Protein

All methylation assays were conducted at 30°C for 1 hour in methyltransferase buffer (50 mM potassium acetate, 50 mM Tris, pH 8.0). 1 μg recombinant NTMT1 was mixed with 1.0 μg full-length recombinant RCC1 and 100 μM SAM and inhibitors added to give the indicated concentrations (2, 10, and 50 μM). Positive controls contained no inhibitor, negative controls contained no NTMT1. One tenth of each reaction was mixed with 5x loading dye and loaded onto a 10% SDS–PAGE gel. Gels were transferred to nitrocellulose membranes using a Biorad Trans-Blot SD Semi-Dry Transfer Cell. Membranes were blocked for 1□h in 5% milk/TBST, then incubated with anti-me3-SPK (1:10,000), anti-me2-SPK (1:5000), or NTMT1 (1:1000) antibodies overnight at 4□°C. They were then washed three times with 1x TBST and incubated with Rabbit IgG-HRP antibody (Jackson ImmunoResearch) for 1□h at room temperature. The membrane was washed three times with 1x TBST and detected using a Biorad ChemiDoc imaging system.

## ASSOCIATED CONTENT

### Supporting Information

The Supporting Information is available free of charge on the ACS Publications website.

Figure S1. MALDI-MS and HPLC spectra of compounds **1**-**24**; Figure S2. IC_50_ curves of compound **5, BM30, DC431**, and **DC432**; Figure S3. Cell permeability evaluation; Table S1. Selectivity study; Table S2. Crystallography data and refinement statistics (PDB ID: 6WH8)

### Accession Codes

The coordinates for the structure of human NTMT1 in complex with BM30 have been deposited under PDB ID 6WH8. Authors will release the atomic coordinates and experimental data upon article publication.

### Author Contributions

The manuscript was written through contributions of all authors. All authors have given approval to the final version of the manuscript.

### Notes

The authors declare no competing financial interest.

## ACKNOWLEDGMENT

The authors acknowledge the support from NIH grants R01GM117275 (RH), 1R01GM127896-01 (NN), 1R01AI127793 (NN), R01GM112721 (CST), and P30 CA023168 (Purdue University Center for Cancer Research). The SGC is a registered charity (number 1097737) that receives funds from AbbVie, Bayer Pharma AG, Boehringer Ingelheim, Canada Foundation for Innovation, Eshelman Institute for Innovation, Genome Canada through Ontario Genomics Institute [OGI-055], Innovative Medicines Initiative (EU/EFPIA) [ULTRA-DD grant no. 115766], Janssen, Merck KGaA, Darmstadt, Germany, MSD, Novartis Pharma AG, Ontario Ministry of Research, Innovation and Science (MRIS), Pfizer, São Paulo Research Foundation-FAPESP, Takeda, and Wellcome. We also thank supports from the Department of Medicinal Chemistry and Molecular Pharmacology (RH) and Department of Biological Sciences (NN) at Purdue University.

## ABBREVIATIONS

NTMT: protein N-terminal methyltransferase
SAM: S-5’-adenosyl-L-methionine
SAH: S-5’-adenosyl-L-homocysteines
SAHH: SAH hydrolase
PKMT: protein lysine methyltransferase
PRMT: protein arginine methyltransferase
rt: room temperature
TFA: trifluoroacetic acid

## Table of Contents

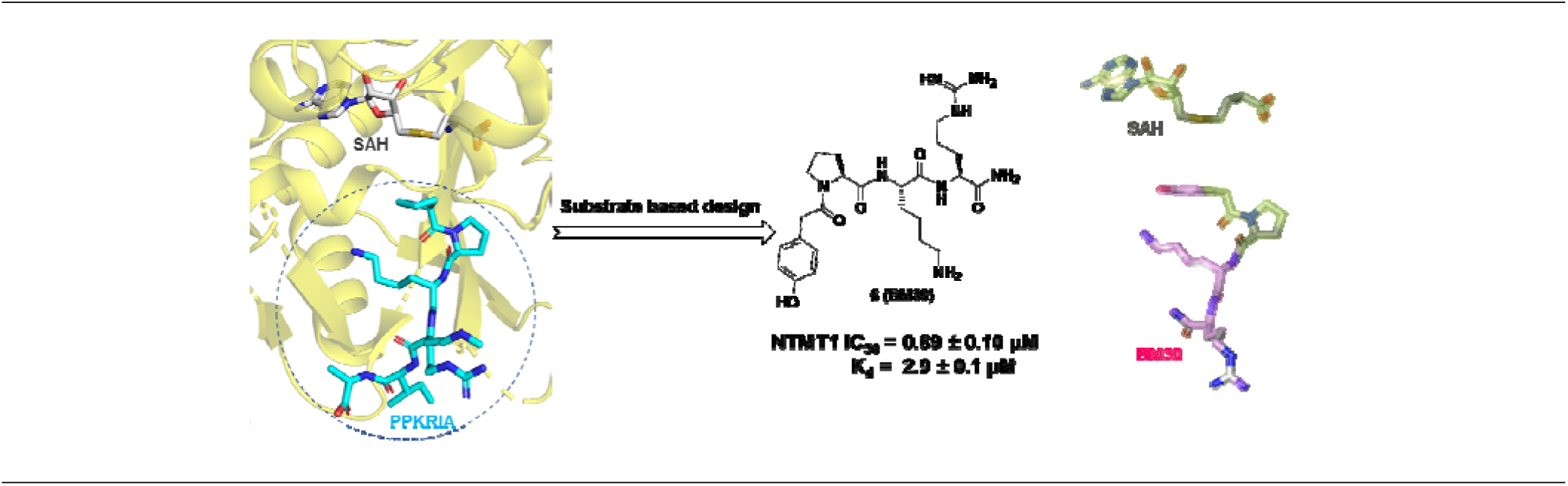

## REFERENCES

(1) Ribich, S.; Harvey, D.; Copeland, R. A. Review Drug Discovery and Chemical Biology of Cancer Epigenetics. Cell Chem. Biol. 2017, 24 (9), 1120–1147.

(2) Scheer, S.; Ackloo, S.; Medina, T. S.; Schapira, M.; Li, F.; Ward, J. A.; Lewis, A. M.; Northrop, J. P.; Richardson, P. L.; Kaniskan, H. Ü.; Shen, Y.; Liu, J.; Smil, D.; McLeod, D.; Zepeda-Velazquez, C. A.; Luo, M.; Jin, J.; Barsyte-Lovejoy, D.; Huber, K. V. M.; De Carvalho, D. D.; Vedadi, M.; Zaph, C.; Brown, P. J.; Arrowsmith, C. H. A Chemical Biology Toolbox to Study Protein Methyltransferases and Epigenetic Signaling. Nat. Commun. 2019, 10 (1), 1–14.

(3) Chan-Penebre, E.; Kuplast, K. G.; Majer, C. R.; Boriack-Sjodin, P. A.; Wigle, T. J.; Johnston, L. D.; Rioux, N.; Munchhof, M. J.; Jin, L.; Jacques, S. L.; West, K. A.; Lingaraj, T.; Stickland, K.; Ribich, S. A.; Raimondi, A.; Scott, M. P.; Waters, N. J.; Pollock, R. M.; Smith, J. J.; Barbash, O.; Pappalardi, M.; Ho, T. F.; Nurse, K.; Oza, K. P.; Gallagher, K. T.; Kruger, R.; Moyer, M. P.; Copeland, R. A.; Chesworth, R.; Duncan, K. W. A Selective Inhibitor of PRMT5 with in Vivo and in Vitro Potency in MCL Models. Nat. Chem. Biol. 2015, 11 (6), 432–437.

(4) Huang, R. Chemical Biology of Protein N-Terminal Methyltransferases. ChemBioChem 2019, 20 (8), 976–984.

(5) Chen, R.; Brosius, J.; Wittmann-Liebold, B. Occurrence of Methylated Amino Acids as N-Termini of Proteins from Escherichia Coli Ribosomes. J. Mol. Biol. 1977, 111 (2), 173–181.

(6) Schaner Tooley, C. E.; Petkowski, J. J.; Muratore-Schroeder, T. L.; Balsbaugh, J. L.; Shabanowitz, J.; Sabat, M.; Minor, W.; Hunt, D. F.; Macara, I. G. NRMT Is an Alpha-N-Methyltransferase That Methylates RCC1 and Retinoblastoma Protein. Nature 2010, 466 (7310), 1125–1128.

(7) Webb, K. J.; Lipson, R. S.; Al-Hadid, Q.; Whitelegge, J. P.; Clarke, S. G. Identification of Protein N-Terminal Methyltransferases in Yeast and Humans. Biochemistry 2010, 49 (25), 5225–5235.

(8) Petkowski, J. J.; Bonsignore, L. A.; Tooley, J. G.; Wilkey, D. W.; Merchant, M. L.; Macara, I. G.; Schaner Tooley, C. E. NRMT2 Is an N-Terminal Monomethylase That Primes for Its Homologue NRMT1. Biochem. J. 2013, 456 (3), 453–462.

(9) Dong, C.; Mao, Y.; Tempel, W.; Qin, S.; Li, L.; Loppnau, P.; Huang, R.; Min, J. Structural Basis for Substrate Recognition by the Human N-Terminal Methyltransferase 1. Genes Dev. 2015, 29 (22), 2343–2348.

(10) Wu, R.; Yue, Y.; Zheng, X.; Li, H. Molecular Basis for Histone N-Terminal Methylation by NRMT1. Genes Dev. 2015, 29 (22), 2337–2342.

(11) Dong, C.; Dong, G.; Li, L.; Zhu, L.; Tempel, W.; Liu, Y.; Huang, R.; Min, J. An Asparagine/Glycine Switch Governs Product Specificity of Human N-Terminal Methyltransferase NTMT2. Commun. Biol. 2018, 1 (1), 183.

(12) Jakobsson, M. E.; Malecki, J. M.; Halabelian, L.; Nilges, B. S.; Pinto, R.; Kudithipudi, S.; Munk, S.; Davydova, E.; Zuhairi, F. R.; Arrowsmith, C. H.; Jeltsch, A.; Leidel, S. A.; Olsen, J. V.; Falnes, P. Ø. The Dual Methyltransferase METTL13 Targets N Terminus and Lys55 of EEF1A and Modulates Codon-Specific Translation Rates. Nat. Commun. 2018, 9 (1), 3411.

(13) Liu, S.; Hausmann, S.; Carlson, S. M.; Fuentes, M. E.; Francis, J. W.; Pillai, R.; Lofgren, S. M.; Hulea, L.; Tandoc, K.; Lu, J.; Li, A.; Nguyen, N. D.; Caporicci, M.; Kim, M. P.; Maitra, A.; Wang, H.; Wistuba, I. I.; Porco, J. A. Jr.; Bassik, M. C.; Elias, J. E.; Song, J.; Topisirovic, I.; Van Rechem, C.; Mazur, P. K.; Gozani, O. METTL13 Methylation of EEF1A Increases Translational Output to Promote Tumorigenesis. Cell 2019, 176 (3), 491–504.e21.

(14) Minoru Nomoto, Yoshimasa Kyogoku, and K. I. N-Trimethylalanine, a Novel Bloacked N-Terminal Residue of Tetrahymena Histone H2B. J. Biochem 1982, 92 (5), 1675–1678.

(15) Sathyan, K. M.; Fachinetti, D.; Foltz, D. R. α-Amino Trimethylation of CENP-A by NRMT Is Required for Full Recruitment of the Centromere. Nat. Commun. 2017, 8, 14678.

(16) Chen, T.; Muratore, T. L.; Schaner Tooley, C. E.; Shabanowitz, J.; Hunt, D. F.; Macara, I. G. N-Terminal Alpha-Methylation of RCC1 Is Necessary for Stable Chromatin Association and Normal Mitosis. Nat. Cell Biol. 2007, 9 (5), 596–603.

(17) Cai, Q.; Fu, L.; Wang, Z.; Gan, N.; Dai, X.; Wang, Y. α-N-Methylation of Damaged DNA-Binding Protein 2 (DDB2) and Its Function in Nucleotide Excision Repair. J. Biol. Chem. 2014, 289 (23), 16046–16056.

(18) Dai, X.; Rulten, S. L.; You, C.; Caldecott, K. W.; Wang, Y. Identification and Functional Characterizations of N-Terminal Alpha-N-Methylation and Phosphorylation of Serine 461 in Human Poly(ADP-ribose) Polymerase 3. J Proteome Res 2015, 14(6), 2575–2582.

(19) Shields, K. M.; Tooley, J. G.; Petkowski, J. J.; Wilkey, D. W.; Garbett, N. C.; Merchant, M. L.; Cheng, A.; Schaner Tooley, C. E. Select Human Cancer Mutants of NRMT1 Alter Its Catalytic Activity and Decrease N-Terminal Trimethylation. Protein Sci. 2017, 26 (8), 1639–1652.

(20) Bonsignore, L. A.; Tooley, J. G.; Van Hoose, P. M.; Wang, E.; Cheng, A.; Cole, M. P.; Schaner Tooley, C. E. NRMT1 Knockout Mice Exhibit Phenotypes Associated with Impaired DNA Repair and Premature Aging. Mech. Ageing Dev. 2015, 146, 42–52.

(21) Hitakomate, E.; Hood, F. E.; Sanderson, H. S.; Clarke, P. R. The Methylated N-Terminal Tail of RCC1 Is Required for Stabilisation of Its Interaction with Chromatin by Ran in Live Cells. BMC Cell Biol. 2010, 11, 43.

(22) Richardson, S. L.; Mao, Y.; Zhang, G.; Hanjra, P.; Peterson, D. L.; Huang, R. Kinetic Mechanism of Protein N-Terminal Methyltransferase 1. J. Biol. Chem. 2015, 290 (18), 11601–11610.

(23) Zhang, G.; Richardson, S. L.; Mao, Y.; Huang, R. Design, Synthesis, and Kinetic Analysis of Potent Protein N-Terminal Methyltransferase 1 Inhibitors. Org. Biomol. Chem. 2015, 13 (14), 4149–4154.

(24) Zhang, G.; Huang, R. Facile Synthesis of SAM-Peptide Conjugates through Alkyl Linkers Targeting Protein N-Terminal Methyltransferase 1. RSC Adv. 2016, 6, 6768–6771.

(25) Chen, D.; Dong, G.; Noinaj, N.; Huang, R. Discovery of Bisubstrate Inhibitors for Protein N-Terminal Methyltransferase 1. J. Med. Chem. 2019, 62 (7), 3773–3779.

(26) Petkowski, J. J.; Schaner Tooley, C. E.; Anderson, L. C.; Shumilin, I. A; Balsbaugh, J. L.; Shabanowitz, J.; Hunt, D. F.; Minor, W.; Macara, I. G. Substrate Specificity of Mammalian N-Terminal α-Amino Methyltransferase. Biochemistry 2012, 51(30), 5942–5950.

(27) Richardson, S. L.; Hanjra, P.; Zhang, G.; Mackie, B. D.; Peterson, D. L.; Direct, Ratiometric, and Quantitative MALDI-MS Assay for Protein Methyltransferases and Acetyltransferases. Anal. Biochem. 2015, 478, 59–64.

(28) De Poli, M.; Moretto, A.; Crisma, M.; Peggion, C.; Formaggio, F.; Kaptein, B.; Broxterman, Q. B.; Toniolo, C. Is the Backbone Conformation of Cα-Methyl Proline Restricted to a Single Region? Chem. – A Eur. J. 2009, 15 (32), 8015–8025.

(29) Chen, D.; Li, L.; Diaz, K.; Iyamu, I. D.; Yadav, R.; Noinaj, N.; Huang, R. Novel Propargyl-Linked Bisubstrate Analogues as Tight-Binding Inhibitors for Nicotinamide N-Methyltransferase. J. Med. Chem. 2019, 62 (23), 10783–10797.

(30) Qian, Z.; Liu, T.; Liu, Y.-Y. Y.; Briesewitz, R.; Barrios, A. M.; Jhiang, S. M.; Pei, D. Efficient Delivery of Cyclic Peptides into Mammalian Cells with Short Sequence Motifs. ACS Chem. Biol. 2013, 8 (2), 423–431.

